# Phloem transport limitation in Huanglongbing affected sweet orange is dependent on phloem-limited bacteria and callose

**DOI:** 10.1101/2021.07.07.451171

**Authors:** Stacy Welker, Myrtho Pierre, James P. Santiago, Manjul Dutt, Christopher Vincent, Amit Levy

## Abstract

Huanglongbing (HLB), caused by *Candidatus* Liberibacter asiaticus (CLas), is a phloem-limited disease which disrupts citrus production in affected areas. In HLB-affected plants, phloem sieve plate pores accumulate callose, and leaf carbohydrate export is reduced. However, whether HLB causes a reduction in carbohydrate phloem translocation speed, and the quantitative relationships among callose, CLas population, and phloem translocation are still unknown. In this work, a procedure was developed to concurrently measure sugar transport, callose deposition, and relative pathogen population at different locations throughout the stem. Increasing quantities of CLas genetic material were positively correlated with quantity and density of callose deposits, and negatively correlated with phloem translocation speed. Callose deposit quantity was site- and rootstock dependent, and were negatively correlated with phloem translocation speed, suggesting a localized relationship. Remarkably, callose accumulation and phloem translocation disruption in the scion was dependent on rootstock genotype. Regression results suggested that the interaction of Ct values and number of phloem callose depositions, but not their size or density, explained the effects on translocation speed. Sucrose, starch, and sink ^14^C label allocation data support the interpretation of a transport pathway limitation by CLas infection. This work shows that the interaction of local accumulation of callose and CLas affect phloem transport. Further, the extent of this accumulation is attenuated by the rootstock and provides important information about the disease mechanism of phloem-inhabiting bacteria. Together, these results constitute the first example of a demonstrated transport limitation of phloem function by a microbial infection.

## Introduction

Huanglongbing (HLB) is a disease of citrus that has severely affected commercial production worldwide (Graça et al. 2016). HLB is putatively caused by several Gram-negative bacterial species, but the most economically significant among these is *Candidatus* ‘Liberibacter asiaticus’ (CLas; Halbert and Manjunath 2004). CLas is vectored by the Asian citrus psyllid, *Diaphorina citri* (Kuwayama). HLB makes citrus production less profitable because growers must invest more in chemical inputs and replacement trees to maintain acceptable yield levels (Farnsworth and Grogan 2014). In Florida, the United States’ historically most productive citrus-producing region, annual fruit yield has fallen by two-thirds, nearly 10 million tons, since HLB was first detected in 2005 (USDA, 2019).

HLB causes symptoms in citrus that resemble nutrient deficiency (Gabriel et al. 2020). Asymmetric chlorosis, often called “blotchy mottle,” can be observed in the leaves. Over a period of years, mature branches die back, causing a loss of canopy volume. New shoots grow in with a characteristic severe chlorosis. Infected trees have a poorly developed fibrous root system (Kumar et al. 2018). The disease also causes fruit drop and malformation. Histologically, the most apparent symptoms include starch accumulation in the leaves, callose deposits blocking the sieve tube pores, and eventually phloem necrosis and collapse (Kim et al. 2009; Achor et al. 2010; Narciso and Exteberria 2015; Deng et al. 2019; Achor et al. 2020).

Previous work has suggested that the symptoms observed in HLB-infected trees can also be induced with stem girdling (Cimò et al. 2013). The phloem collapse, which is observed in HLB infection, may replicate the effect of girdling, leading to systemic phloem translocation dysfunction. The disease may cause a metabolic imbalance over time, as sink tissues become starved by a poor supply of carbon compounds that are needed for normal development. Several lines of evidence support this concept of the HLB disease process. Large structural roots, normally responsible for starch storage, were found to be starch-deficient in HLB-infected citrus trees (Aritua et al. 2013; Kumar et al. 2018). Fruit from CLas-positive trees contains markedly less sugar (Rosales and Burns 2011). Massive quantities of starch can be found in the mature leaves in trees with advanced HLB (Deng et al. 2019). However, other studies suggest that in the roots only the presence of CLas, in conjunction with a non-callose phloem plugging mechanism, was sufficient for drastic damage to the host to occur (Johnson et al. 2014; Achor et al. 2020). Further research is required to resolve the connection between callose plugging, phloem carbon transport dysfunction, and the presence of CLas bacteria.

Historically, it has been difficult to measure the transport of carbon through phloem tissue. Physical handling disrupts normal phloem tissue function during experiments. However, a method of using x-ray detectors to detect ^14^C movement following a pulse from photosynthesized ^14^CO_2_ allows the direct measurement of phloem flow rates in living plants (Epron et al. 2012; Vincent et al. 2019). ^14^CO_2_ gas infusions have been used to demonstrate photoassimilate transport dysfunction in HLB-infected trees in previous work, however, only the leaf ^14^C export was measured (Koh et al. 2012). By tracking the speed of phloem translocation through the stem, the influence of HLB-induced callose plugging on carbohydrate (CHO) transport can be better studied.

Callose is a gelatinous, β-1,3-linked polysaccharide of glucose which is used by plants for a variety of structural and developmental functions (Piršelová and Matušíková 2013). It is produced by transmembrane proteins called callose synthases (CalS) or glucan-synthase-like (GSL) proteins. In *Arabidopsis*, 12 CalS have been described, many of which have homologs in *Citrus* (Verma and Hong 2001; Granato et al. 2019). One function for which plants use callose is to regulate the aperture size of the plasmodesmata and sieve pores in the phloem (Amsbury et al. 2018). Experimental evidence shows that this callose production in the phloem contributes to immunity against many different types of pathogens (Enrique et al. 2011; Ellinger et al. 2013; Fernández-Crespo et al. 2017). Additionally, phloem callose decreases the transport of photoassimilates and auxins, while treatments that stimulate breakdown of sieve plate callose increase movement of fluorescein through the sieve tubes (Webster and Currier 1965; McNairn and Currier 1968; Hollis and Tepper 1971; McNairn 1972; Aloni et al. 1991; Maeda et al. 2006). Multiple groups have produced electron micrographs which show CLas bacteria interacting with host cell membrane and passing through the sieve pores when callose was not present (Kim et al. 2009; Achor et al. 2020). It is thought that the presence of CLas induces callose formation as a defense response. Previously, callose deposits in HLB-infected citrus phloem have been observed by staining the leaf petiole with aniline blue, taking images of thin sections, and either manually counting callose deposits or measuring the number of illuminated pixels (Kim et al. 2009; Boava et al. 2017).

The objective of this work is to better-understand the link between callose formation, CLas, and sugar translocation in the phloem. For this, a high-throughput method for counting and assessing the density of callose deposits in confocal imagery, in combination with a noninvasive measurement of phloem transport speed, and qPCR quantification of CLas were applied to the citrus phloem for the first time in the same tissues along the stem.

Measurements were carried out in healthy and infected sweet orange scions grafted on two different rootstocks. We provide evidence for the correlation of the presence of CLas bacteria and phloem callose plugging with systemic phloem dysfunction.

## Materials and methods

### Plant materials and growth conditions

This experiment used a randomized complete block design. Plants consisting of a ‘Hamlin’ sweet orange (*Citrus sinensis* L.) scion on either ‘Cleopatra’ mandarin (*C. reticulata* Blanco) or ‘X-639’ (*Poncirus trifoliata* L. x *C. reticulata* ‘Cleopatra’) rootstock were kept in a greenhouse with a natural day/night cycle. Each rootstock group consisted of 6 plants, half of which were infected with HLB through infected psyllids during Aug. 2017. This created four treatment groups with 3 plants each: HLB+/X-639, HLB-/X-639, HLB+/Cleo, and HLB-/Cleo. To ensure that phloem translocation was occurring only from the mature leaves to the roots, trees were pruned to promote new shoot growth, and measurements were taken on the woody stem after the maturation of all new shoots.

### Assessment of phloem translocation speed

The phloem transport speed of the trees in this study was measured using previously described methods (Vincent et al. 2019). Trees were individually placed in a radioisotope-rated fume hood for the duration of the pulse-chase period. Inside the hood, plants were given a 12-hour light cycle using an LED grow light, with a leaf-level photosynthetic photon flux density of approximately 1000 µmol m^-2^ s^-1^. Four x-ray photomultiplier tubes (St. Gobain, Malvern, PA) were positioned along the stem and held in place with clamps and stands, allowing for the translocation speed to be assessed for the 3 regions between each detector (Figure 1). The distance between the detectors was measured for each stem.

**Figure 1.**
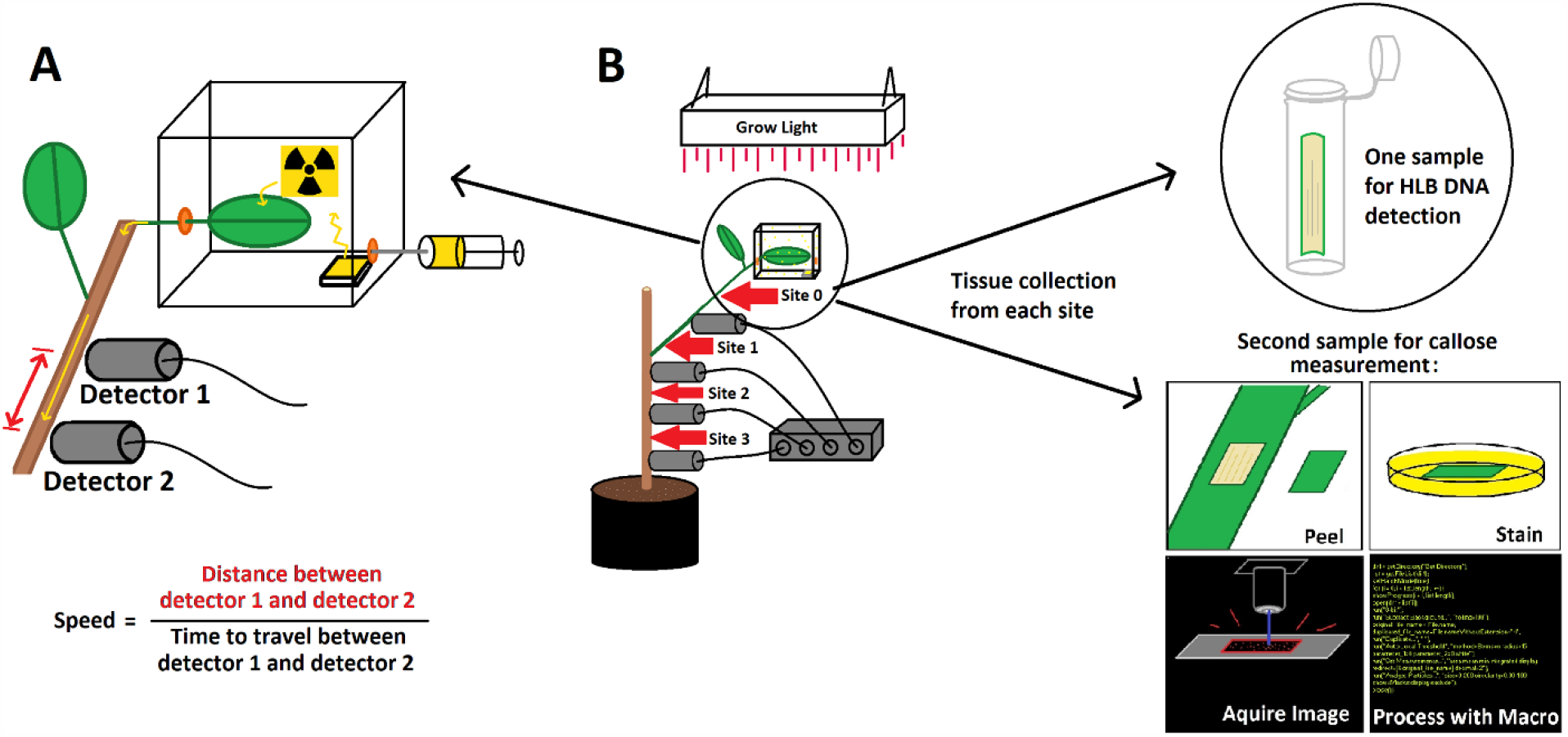
Detail of experiment set-up for infusion of a ^14^C tracer into live citrus plants. **A:** Chamber and detector set-up. Attached leaves were sealed in an air-tight chamber with access to light. Through a rubber septum in the chamber, radiolabeled bicarbonate (Na^14^CO_3_) and citric acid (C_6_H_8_O_7_) are injected into a cuvette, releasing ^14^CO_2_ gas. The ^14^C is incorporated into the photosynthetic reactions of the plant, creating radiolabeled sugars which can be tracked through the phloem using x-ray detectors. After the experiment, the phloem translocation speed is calculated with a simple formula as depicted. **B:** Overview of experiment set-up. Four x-ray photomultiplier tubes were positioned along the stem of each tree. Stem phloem tissue was sampled at sites 0-3 (red arrows). Half of each stem sample was fixed and stained to measure callose. DNA was extracted from the other half to assess the relative quantity of CLas genetic material.

A single leaf at the distal end of a branch, still attached to the tree by the petiole, was sealed inside an air-tight cuvette (Figure 1). Light passed through the plexiglass lid with approximately 90% transmission so that photosynthesis could occur in the leaf as normal. Trees were kept in the hood for 24 hours prior to the assay to allow any callose induced by movement during transportation to be degraded. Following the waiting period, 100 μL of liquid Na^14^CO_3_ (100 µCi) was injected into a receptacle through a septum in the cuvette. Immediately afterwards, 500 μL of 1M citric acid solution was also injected into the cuvette. A 30-minute pulse period elapsed, after which the container was opened and the ^14^CO_2_ gas was vented by the fume hood. The *Bremsstrahlung* radiation was detected by the photomultiplier tubes and logged each minute with an M4612 12-channel counter (Ludlum Measurements, Sweetwater, TX, USA).

### Quantification of phloem callose deposit number and density

This method was adapted from a protocol to measure callose in the *Arabidopsis* plasmodesmata (Zavaliev and Epel 2015). To obtain samples of the phloem, a scalpel was used to collect peels of stem bark tissue approximately 1.5 to 2.5 cm in length and 0.5 to 0.75 cm in width from each tree. The stem peels were taken from four sites at different sections of stem on each tree (Figure 1). The sampling sites corresponded with the location of each ^14^C detector from the phloem translocation speed assay. To prevent spurious measurement of callose deposits caused by wounding, the stem peels were placed immediately after collection into 2 mL Eppendorf tubes containing 85% ethanol on ice. The stem peels were de-stained in the ethanol solution overnight at room temperature. The peels were rehydrated for 1 hour in a 0.01% Tween-20 solution. Afterwards, the peels were stained for 1 hour with an aniline blue solution prepared according to a protocol developed previously (Levy et al. 2007). The peels were placed on glass slides with a drop of 0.01% Tween-20 solution.

Images were taken with a Leica SP8 confocal microscope using methods modified from previous work (Zavaliev and Epel 2015). A 405 nm diode laser was used for excitation and a 475-525 nm band-pass filter was used for emission detection. The laser power, gain, and offset were optimized to minimize background noise and oversaturation and the same settings were used for every image. Five images were taken each from 3 peels from each site on the trees. Images were taken randomly from the central portion of each peel, ensuring that no images overlapped. Callose deposit count per 10X field and integrated density values of callose deposits were collected using Fiji/ImageJ software. Integrated density is the product of each deposit’s area and mean grey value. It is reported in relative fluorescent units (RFU). A macro was modified from previous work to collect data in a standardized, automated manner (Zavaliev and Epel 2015). The macro settings were optimized to identify and measure citrus phloem callose deposits.

### Total nucleic acid extraction

Stem phloem samples from each plant were collected from the same sites as the callose peel collection (Figure 1). Leaf petiole tissue, traditionally thought to be the sample most likely to display positive qPCR results if the pathogen is present, was also collected from each plant. Each sample consisted of approximately 100 mg of tissue. DNA was extracted from the samples in a manner similar to previous work (DePaulo and Powell 1995; Irey et al. 2006), using a modified DNA extraction buffer that contained 0.1 M NaCl, 10 mM EDTA, 50 mM Tris at pH 9.0, and 10 mM DTT. The DTT was added immediately before use. The DNA concentration of each sample was checked and standardized to 100 ng/μL.

### HLB 16S genetic material expression analysis

Quantitative real-time PCR was performed to detect HLB genetic material in total nucleic acid samples that extracted as described above. The HLBas and HLBr primer and HLB-p-FAM probe sequences were used as described previously (Li et al. 2006). The 10 μL reaction mixture was prepared as follows: 5 μL TaqMan™ Fast Universal PCR Master Mix (Applied Biosystems), 0.2 μL each of forward and reverse primers (0.2 uM), 0.1 μL probe (0.1 uM), 3.5 μL DNAase free water, and 1 μL of DNA (100 ng/μL). For processing the reactions, an Applied Biosystems™ 7500 Real-Time PCR System was used with the following run settings: 94°C for 5 minutes, 40 cycles at 94°C for 10 seconds and 58°C for 40 seconds. The thermocycler’s 7500 Fast system SDS software was used with a manual threshold of 0.2 to record the Ct values. Three technical replicates were performed and averaged to obtain each sample result. Similar to previous work, samples with a Ct value of 32 or less were considered positive for CLas, while samples with a value of 32 or higher were considered negative (Irey et al. 2006). For ease of interpretation, the HLB Ct values were expressed as ΔCt, or the difference between the negative (no-template) control Ct and experimental tree Ct. When expressed this way, a higher ΔCt value indicates that a higher amount of HLB DNA was detected.

### Quantification of starch, sucrose, and relative sink activity

For these assays tissues from the same segments as the callose samples were gathered. Additionally, we sampled unlabeled leaves from the apical and basal portions of the canopy as well as the labeled leaf, and we sampled cortical tissue from structural roots and fibrous roots (diameter <2 mm). These were frozen in liquid nitrogen and ground with a chilled tissue lyser. The ground samples were then lyophilized for 24 hrs. A subsample of 100 mg was used to quantify ^14^C activity using liquid scintillation (Beckman 1400; Beckman Coulter, Brea, CA, USA). Thus, ^14^C activity, sucrose and starch were quantified on the same homogenized tissue samples. Relative sink activity was calculated as the ^14^C activity of each tissue relative to the combined activity of all sink tissues (including non-labeled leaves) measured in the same plant.

Sucrose was extracted from the lyophilized tissue with 300 µL of 3.5% perchloric acid, homogenized, and incubated on ice for 5 min. The slurry was centrifuged for at 18,000 x g for 10 min at 4°C. The pellet was set aside, and the supernatant was transferred to a new tube and neutralizing buffer was added (2 m KOH, 150 mM Hepes, 10 mM KCl) at 0.25 volume of the supernatant to bring up the pH to about 7.0. The mixture was then frozen and thawed twice using liquid nitrogen to precipitate the potassium perchlorate and then centrifuged as described above. The supernatant was transferred to a new tube and assayed for sucrose content. For starch extraction, the tissue pellet was resuspended with 850 µL of double distilled water and centrifuged 2X at 18,000 x g for 10 min at 4°C. After performing 2X and discarding the supernatant, 850 µL of 80% ethanol was added, vortexed, and centrifuged at 18,000 x g for 10 min at 4°C. After the supernatant was discarded, and the pellet was dried in a speed vac concentrator to remove any remaining ethanol. The dried pellet was completely resuspended in 500 µL of 200 mM KOH and then incubated at 95°C for 30 min. The samples were allowed to cool at room temperature after incubation and then 90µL of 1 M acetic acid was added and vortexed. After, 200 µL of 200 mM sodium acetate, pH 5.0 was added and vortexed. Finally, 50 µL of enzyme cocktail (6.6 U amyloglucosidase and 50 U ⍰-amylase) was added, vortexed, and then the samples incubated on a shaker at room temperature for 3 days. After centrifugation, the supernatant was transferred to a new tube and used for starch measurement. Sucrose and starch were measured using a NADPH-linked assay using a plate reader (Soltani et al., 2019). Briefly, samples were mixed with a buffer containing 110 mM Hepes, 488 µM NADP, 488 µM ATP, and 0.4 U glucose 6-phosphate dehydrogenase transferred to a 96-well plate and a stable baseline optical density (OD) was measured at 340 nm. Hexokinase (1 U) and invertase (50 U) were added to each well and the change in OD was recorded. Sucrose and starch were measured as glucose equivalents and calculated using the molar extinction coefficient of NADPH at 340 nm that is 6220 M^-1^ cm^-1^.

### Statistical Analysis

All data gathered from the experiments was analyzed using R (R Core Team 2020). Assumptions for each statistical testing method were checked and if needed, data sets were transformed to meet assumptions as noted. Unless otherwise stated, analyses of variance were performed using the a linear model in base R, including the fixed effects of CLas status, rootstock, and location (sampling site) where relevant, all as fixed effects. Location was nested within plant. Because the design was a completely randomized design, no random effects were included. Differences in mean callose deposit integrated density between treatment groups were analyzed using a one-way analysis of variance (ANOVA) on Box-Cox transformed data. In all models, site was nested within tree. Post-hoc analysis was conducted for the stem site means using Tukey’s HSD test.

To calculate the phloem ^14^C translocation speeds, a logistic regression method, modified from previous work, was used (Vincent et al. 2019). An average time of arrival of the ^14^C tracer in the tree phloem at each photomultiplier tube detector was obtained by fitting a logistic curve to the counts per minute as a function of time. The midpoint of each curve on the x-axis was considered the average time of first arrival of the pulse at the detector. The phloem translocation speed at each point measured along the stem was obtained with the calculation:

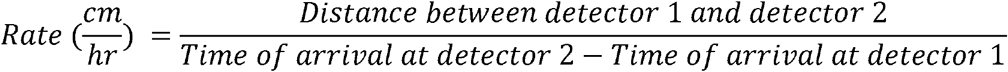

Leaf carbohydrate export was quantified by calculating the pre-labeling baseline, and the post-labeling maximum signal the x-ray detector touching the abaxial side of the labeled leaf. The variable used was the proportion of signal reduction (%) at 8 hrs after the end of the labeling.

The distribution of the callose deposit count data was highly skewed towards zero, meaning that callose deposits were not detected in most images of the phloem. It is likely that the images that were reported as having a count of zero callose deposits did contain some callose, but the authors calibrated the microscope gain and offset settings to avoid saturation and excess noise in the images. This likely caused the most weakly fluorescent callose deposits to not be captured. The count data were also highly dispersed, indicating that callose deposits number varied widely.

For statistical analysis of count data with a highly dispersed distribution and a large number of zeros, a zero-altered negative binomial (ZANB) model, also called a hurdle model, is often the most appropriate (Yang et al. 2017). A related type of analysis, the zero-inflated negative binomial model (ZINB) which accounts for structural and non-structural zeroes, may also have been appropriate. The Akaike information criterion (AIC) was compared between the ZANB and ZINB models for the callose deposit count data and the ZANB was determined to fit better. ZANB models with different independent variables and their interactions were compared with ANOVA tests to determine which was most parsimonious.

The ZANB model for the callose deposit count data was computed with the {pscl} package (Zeileis et al. 2008; Jackman 2017). The model split the data into two groups: zero counts which were analyzed with a logistic regression, and non-zero counts which were analyzed with a negative binomial regression. This allowed us to separately model the factors which influence the odds of callose forming or not and if callose formed, what factors affected the amount of deposits. In both regressions, the effect of the rootstock type, the site on the plant stem where the sample was collected, and the CLas status of the plant were examined as factors in callose formation.

Correlations between mean callose deposit counts per 10x field, mean integrated density, mean phloem translocation speed, and mean CLas ΔCt values were checked using the {GGally} and {hmisc} packages in R (Harrell Jr and Dupont 2020; Schloerke et al. 2020). The means of callose deposit counts per 10x field image were calculated for the measurements from each rootstock, CLas status, and site group within each plant. When parameters exhibited significant correlations, a linear regression was calculated. The assumptions for the model were checked using the Shapiro-Wilk test and by plotting the residuals of the model. For the dataset used in the linear regressions, sites on individual plants which were missing values in any relevant variable were omitted. Two extreme outliers which represented technical errors were dropped from the phloem translocation speed data set.

To determine the most parsimonious model for effects contributing to phloem transport speed, the linear model was developed in negative stepwise fashion. Beginning with the full model, we removed the highest-order non-significant interactions, one at a time. In each step we compared the more complex with the simplified model using a likelihood ratio test to compare models.

## Results

### Phloem translocation speeds and leaf carbohydrate export-rate and allocation varied with CLas status and rootstock type

Using the setup described in Figure 1, we measured the bacterial accumulation, phloem translocation speed, and callose accumulation in three different sites (1-3) along the stem of Hamlin sweet orange scions on either Cleopatra (*Citrus reticulata*) or X-639 (Cleopatra mandarin x *Poncirus trifoliata*) rootstock. CLas DNA was quantified in each stem segment of healthy and infected trees using qPCR. The rootstock type and site on the plant from which the tissue sample was obtained did not have a significant effect on the ΔCt value. The only factor that significantly influenced the ΔCt value was whether the plant was in the CLas+ group (Supplemental Table S1).

For the translocation speed, there was an interaction between rootstock type and CLas status (Supplemental Table S2). In healthy plants, those on X-639 rootstock had a faster translocation speed than those on Cleopatra, but when the plants were challenged with CLas, the slowing effect was much greater in those with X-639 rootstock (Figure 2).

**Figure 2.**
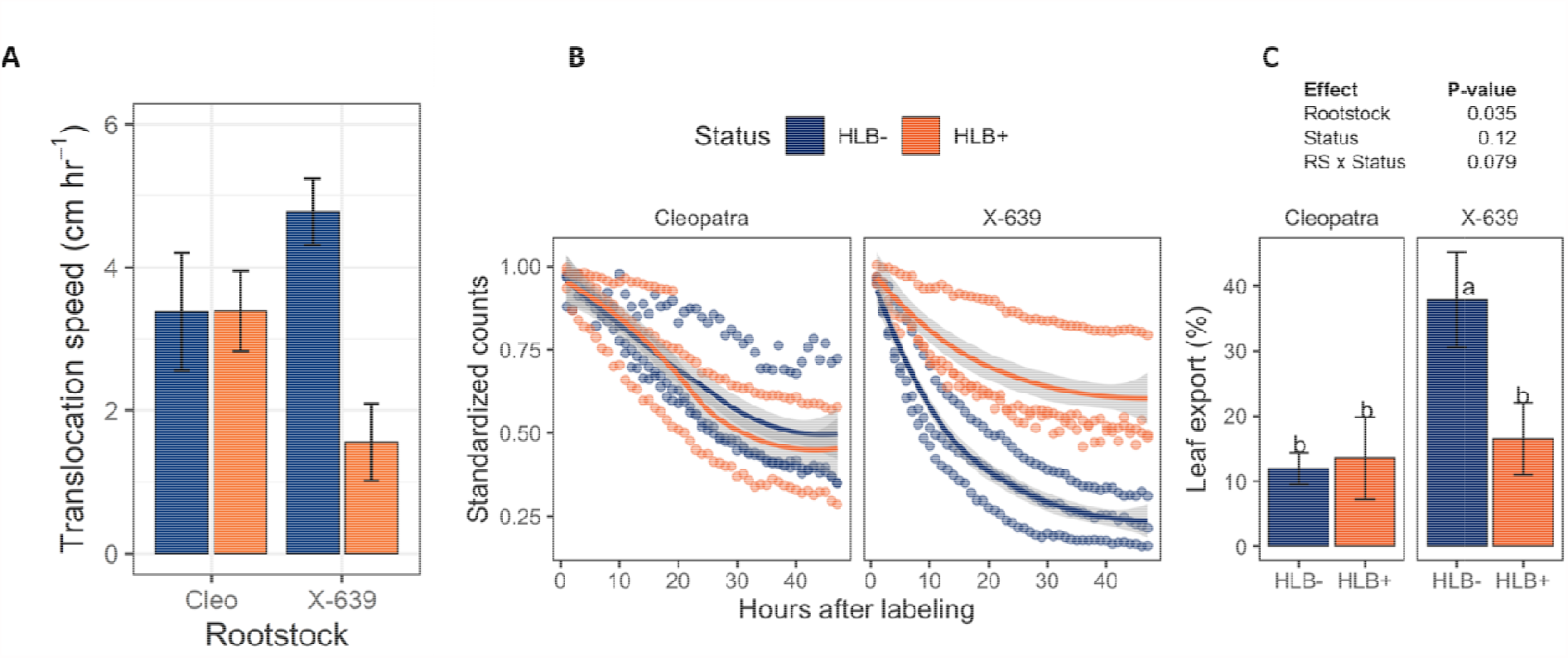
(A) Phloem translocation speeds obtained from Hamlin sweet orange (*Citrus sinensis*) scion on Cleopatra mandarin (*C. reticulata*) or X-639 (*C. reticulata* x *Poncirus trifoliata*) infected or uninfected by *Candidatus* Liberibacter asiaticus and treated with a ^14^CO_2_ pulse-label. Bars represent means and error bars represent standard error, n = 9 for all groups except X-639/CLas+, where n = 6. All contrasts are significant at *P*< 0.05 according to analysis of variance results. (B-C) Export of photosynthetically fixed ^14^C label from labeled leaves B: decline in standardized counts over 48 h after labeling. C: analysis of variance and results of the % export after 8 h. Bars show means, and error bars show standard error. Bars labeled with different letters are significantly different at P<0.05 by Fisher’s LSD

Using x-ray detectors in direct contact with the abaxial side of the leaf, we measured the export of carbohydrates as the decline in signal from the labeled. The leaf export-rate pattern was consistent with the phloem translocation speeds. There was a possible interaction of rootstock and CLas status (F=4.02, *P*=0.079) and a significant rootstock effect (F=6.41, *P*=0.035), in which plants on X-639 had a higher rate of leaf export than those on Cleopatra, but CLas infection greatly reduced export of leaves on X-639 but not those on Cleopatra (Figure 2).

To assess allocation and carbohydrate status, we measured sucrose and starch concentrations and ^14^C activity in samples in non-labeled leaves, trunk cortex from the same segments as the transport and callose measurements and structural and fine roots. For leaf sucrose there was an interaction of rootstock and CLas status (F=6.0, *P*=0.04, Supplemental Table S3), but post-hoc analysis found no differences among means at α<0.05 (Supplemental Figure S1). A 3-way interaction among rootstock, CLas status, and location was found for sucrose concentrations in roots (F=4.25, *P*=0.04). CLas infection reduced sucrose concentrations minimally in fibrous roots, while infection decreased sucrose greatly in structural roots of X-639 but not in those of Cleopatra (Figure 3). There was a possible status x location interaction for starch concentration (F=3.4, *P*=0.067; Supplemental Table S3). In this interaction, CLas infection decreased starch in fibrous roots, but not in structural roots (Supplemental Figure S2). There was an interaction of infection status x rootstock for proportional ^14^C sink activity (F=8.39, *P*=0.01). In Cleopatra, there was no effect of CLas status on label activity of roots, but CLas infection significantly increased the label activity in roots of X-639 (Figure 3).

**Figure 3.**
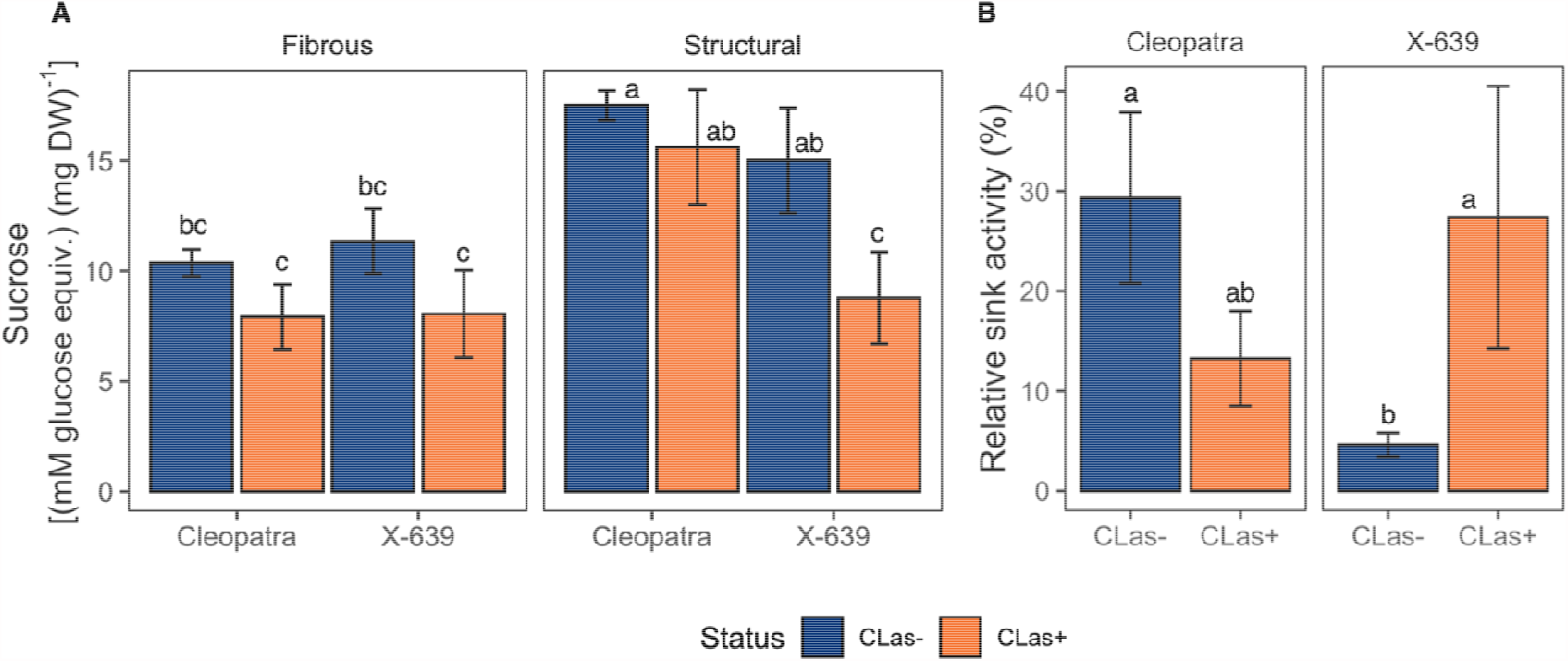
(A) Sucrose concentrations in fibrous roots and structural root cortical tissues of Cleopatra mandarin (*Citrus reticulata*) or X-639 (Cleopatra x *Poncirus trifoliata*) infected or uninfected by *Candidatus* Liberibacter asiaticus, all with Hamlin sweet orange (*C. sinensis*) scion. Bars show means, and error bars show standard error. Bars labeled with different letters are significantly different at P<0.05 by Fisher’s LSD. (B) Relative ^14^C sink activity of fibrous roots and structural root cortical tissues. Samples were gathered 3 days after photosynthetically labeling an apical leaf. Bars show means and error bars show standard error. Bars labeled with different letters are significantly different at P<0.05 by Fisher’s LSD.

### Number and density of callose deposits are influenced by site on plant stem, rootstock type, and CLas status

Next, we measured the average number and the integrated density of callose deposits in the phloem of healthy and infected plants at each stem site (Figure S3). The results of the ZANB analysis on the callose deposit counts per 10x field image are shown in Figure 4 and Supplemental Table S4. Two 3-way combinations of factors greatly affected the odds of callose formation. Plants had higher likelihood of forming callose at sites 0 and 3 if they were CLas+ and on Cleopatra rootstock. There were no 3-way combinations that significantly influenced the amount of callose that formed when the deposit count was greater than zero, however, several two-way interactions were significant. CLas+ plants on X-639 rootstock produced significantly more callose than CLas+ on Cleopatra. Callose deposits were significantly more numerous at sites 2 and 3 with Cleopatra rootstock. When plants were CLas-positive, they were significantly less likely to form any callose at site 3. When callose did form in infected plants, deposits were significantly less numerous at site 3. The callose deposit integrated density measurements for each treatment group were compared using an ANOVA (Supplemental Table S5). The interaction between rootstock, CLas status, and site on plant had a significant effect on the callose deposit integrated density.

**Figure 4.**
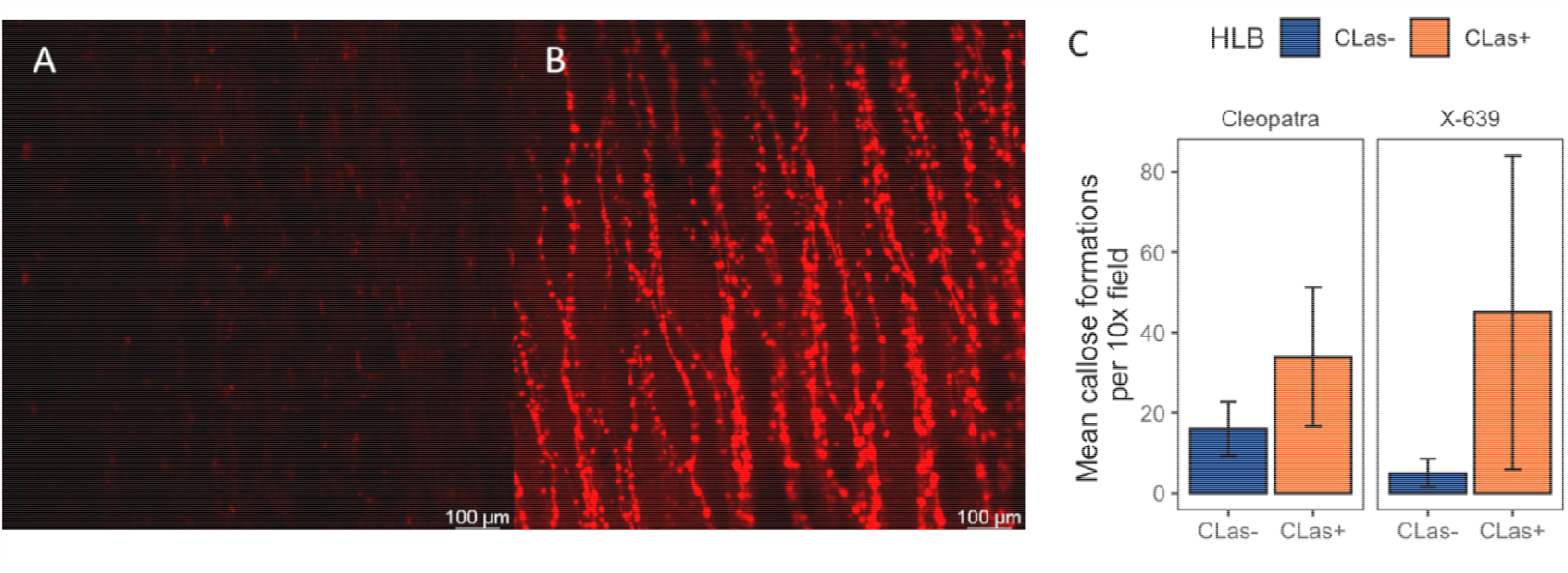
(A-B) Representative images of citrus stem phloem tissue samples. (A) A low-callose image from an uninfected plant. (B) A high-callose image from an HLB-infected plant. (C) Mean number of callose deposits detected per 10x field in confocal micrographs of citrus phloem. One 10x field image captures an area of phloem tissue which is approximately 1.16 mm^2^. Error bars represent standard error, number over each error bar represents n. All contrasts are significantly different at *P*= < 0.05 by Fisher’s LSD.

### Quantity of CLas genetic material and callose deposit count have the greatest influence on phloem translocation speed

Pearson’s correlation coefficients were calculated among all variables (Table 1). Phloem translocation speed had significant negative correlations with both mean callose deposit count and mean ΔCt. Mean ΔCt had positive correlations with mean callose deposit count. The correlations were used to guide further exploration of the datasets with linear regression between the variables that were most correlated (Figure 5). ΔCt affected callose deposit number. As ΔCt values increase, indicating a larger quantity of CLas genetic material, callose deposits became more numerous. Furthermore, callose deposit count affected phloem translocation speed. Higher counts of callose deposits were associated with slower phloem translocation speeds.

**Table 1.**
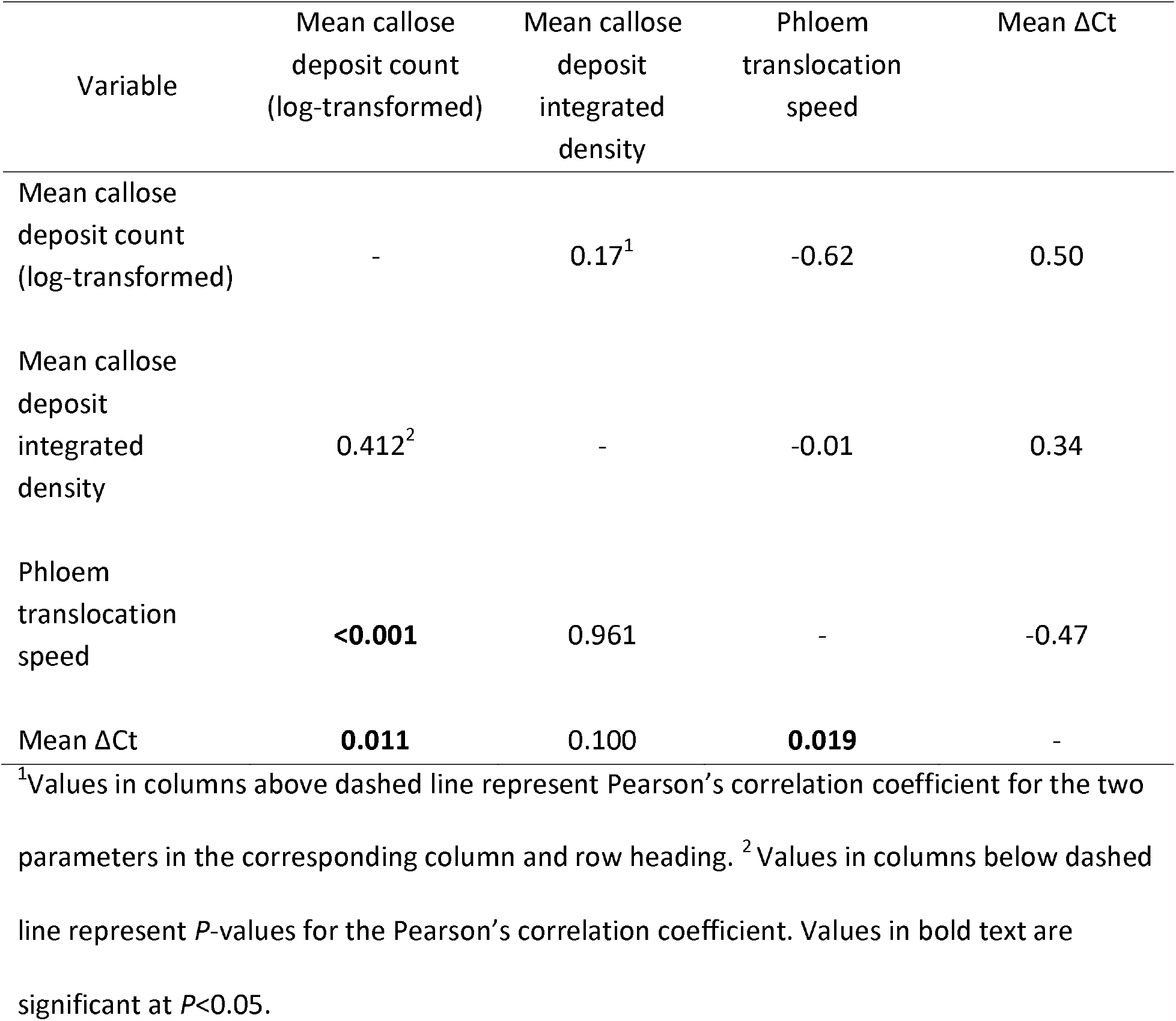
Summary of correlations among the mean callose deposit count per 10x field image, mean callose deposit integrated density (RFU), translocation speed (cm hr^-1^), and ΔCt values. All parameters were measured in the phloem of the same stem segments of CLas+ and CLas-Hamlin sweet orange on either X-639 or Cleopatra rootstock.

**Figure 5.**
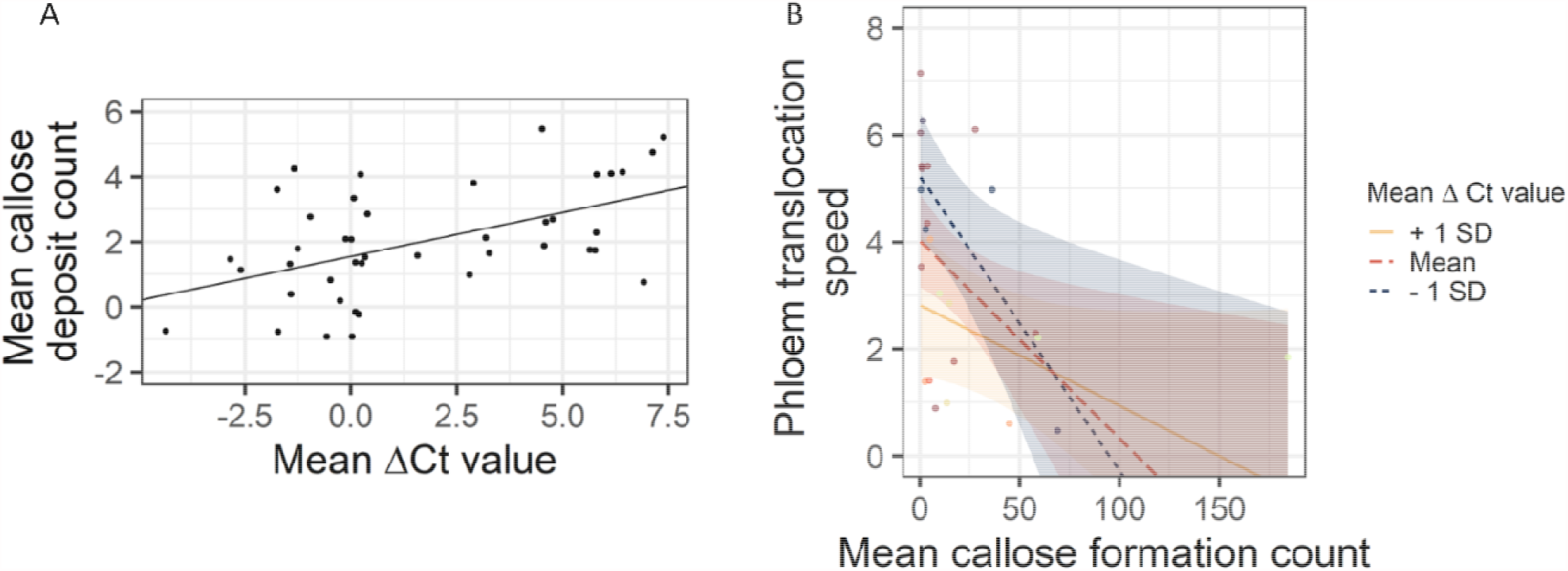
(A) Linear regression of the log-transformed mean callose deposit count per 10x field image by mean ΔCt value. Adjusted R^2^= 0.24, P <0.001. (B) Linear regression of phloem translocation speed (cm hr^-1^) by log-transformed mean callose deposit count per 10x field image. Adjusted R^2^= 0.24, P= 0.004. Red represents the best-fit regression line. Orange and blue lines represent translocation speed plotted as a function of mean callose deposit counts at a different level of ΔCt value to illustrate the relationship between phloem translocation speed, mean callose deposit counts, and ΔCt value. Shaded area represents the 95% confidence interval.

Based on the correlations between the factors, a multiple linear regression model was developed to determine the most influential factors that affected phloem transportation speed (Table 2). Although the model initially included infection status and rootstock variety, these factors were eliminated in the stepwise process, indicating that ΔCt value and mean callose deposit count per 10x field image explain the variation due to rootstock and HLB treatment. According to the coefficients, ΔCt value had the greatest negative effect on the translocation speed, although callose deposit count also contributes to slower speeds. In our dataset, for every additional callose deposit per 10x field, the phloem speed decreased by 0.04 cm hr^-1^. Finally, there was an interaction between ΔCt value and callose deposit count. This interaction had a slightly positive coefficient. Figure 5 illustrates how the relationship between translocation speeds and callose deposit counts changed at different ΔCt values. At negative values of mean ΔCt and those that are close to zero, the best-fit line between mean callose deposit count and mean phloem translocation speed has a steep negative slope. However, at high values of mean ΔCt, the slope of this line flattens, indicating a less dramatic decrease in phloem speed with increasing callose but a lower translocation speed at the x-intercept (i.e., a lower speed in the absence of callose).

**Table 2.** Linear model which describes the most significant predictors of mean phloem translocation speed.

| Effect | Estimate | Standard Error | t-value | P-value |
| --- | --- | --- | --- | --- |
| (Intercept) | 4.50 | 0.44 | 10.28 | <0.001 |
| Mean ΔCt value | -0.42 | 0.15 | -2.76 | **0.012** |
| Mean callose deposit count | -0.04 | 0.02 | -2.56 | **0.018** |
| Mean ΔCt value x Mean callose deposit count | 0.01 | <0.01 | 2.31 | **0.031** |

Note: Degrees of freedom= 21, adjusted $R^2$ = 0.32.

## Discussion

The speed reported here (2.5-5.2 cm hr^-1^ for CLas-negative trees) place *Citrus* outside the typical range for angiosperm trees. Studies of angiosperm phloem-transport speed have ranged from 13-122 cm hr^-1^ using a variety of methods to estimate speed (Liesche et al. 2013). These studies have used temperate woody species, and most measures have been greater than 30 cm hr^-1^. The citrus phloem speeds presented here are within the range of those reported for gymnosperms, which range from 2.7-100 cm hr^-1^, but with most measures between 5 and 20 cm hr^-1^ (Liesche et al. 2013). These low speeds are in agreement with sieve plate pores ultrastructure in citrus where pore diameters are smaller than many other plant species reported (Esau and Cheadle 1959; Achor et al. 2020). One important difference between citrus and other angiosperms is its evergreen growth habit. Interspecies phloem speeds have not been well explored, but differences between deciduous and evergreen species may represent differences in carbon budget requirements among canopies with different leaf turnover rates.

Although studies of phloem transport speed have not typically addressed phloem dysfunction, de Schepper et al. (2013) observed the speed and spatial path of *Quercus rubor* transport before and after the imposition of a partial girdle. They observed that girdling one-half of the stem induced slower transport on the same side of the stem as the girdle but accelerated transport on the opposite side. Below the girdle, transport was not slowed. A rerouting of transport around the girdle was proposed as the cause of the relative maintenance of transport.

Our experimental results confirm previous studies that suggested callose is produced by infected citrus as a response to CLas presence and as a response to limit its movement through the phloem (Folimonova et al. 2009; Koh et al. 2012; Boava et al. 2017; Deng et al. 2019; Achor et al. 2020). Like the re-routing that occurs because of girdling, if phloem sieve tubes are blocked with callose in CLas-infected plants, then phloem transport must proceed at a reduced rate through alternate routes unless no additional alternative routes exist (Figure 6). Other studies have demonstrated that, in response to this situation, the plant hyperstimulates its vascular cambium leading to phloem regeneration, especially in tolerant varieties (Deng et al. 2019). This phenomenon may explain the tolerance of these varieties.

**Figure 6.**
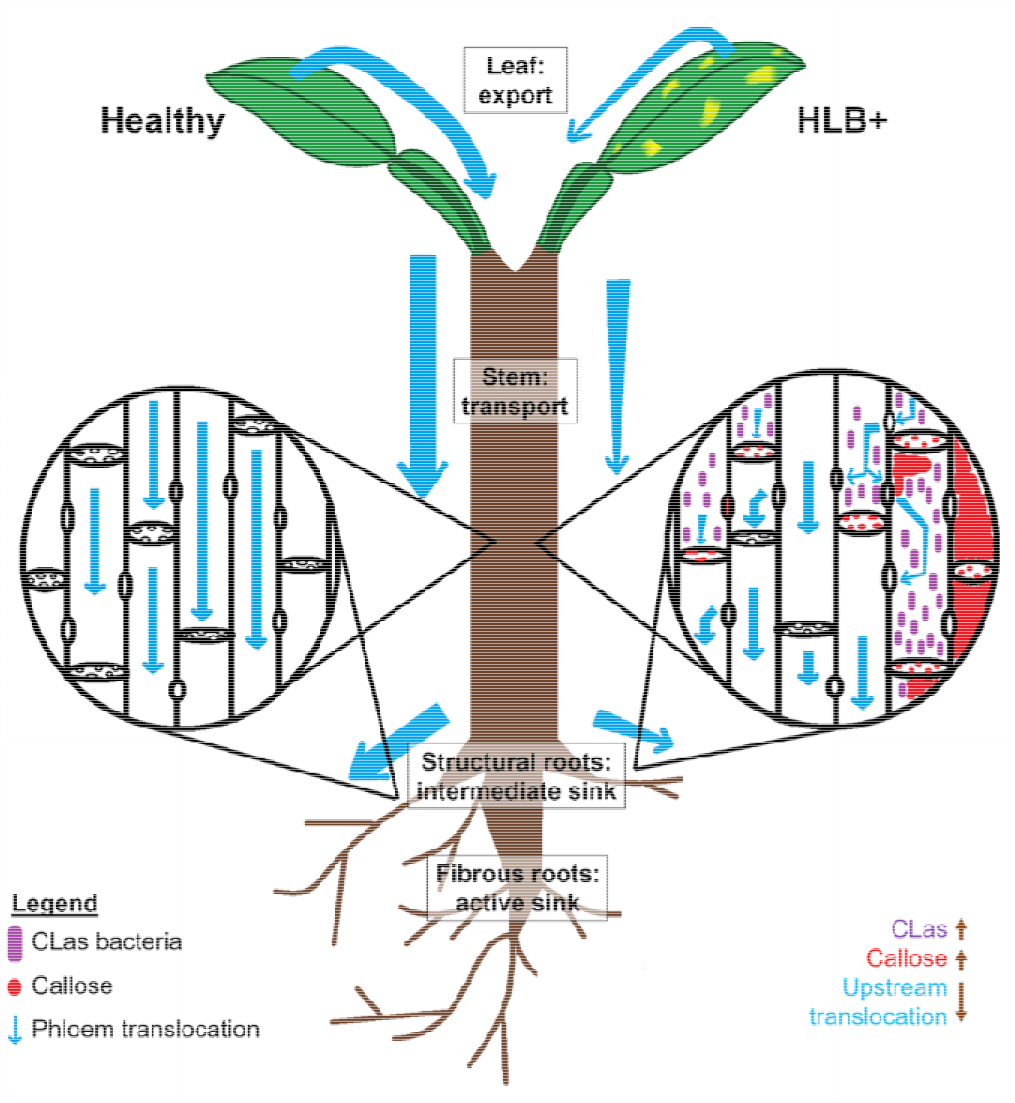
Proposed model of phloem transport differences in HLB-infected and healthy plants. The presence of CLas is associated with increased callose formation, and together these two factors are associated with a decrease in stem phloem carbohydrate transport speed. CLas bacteria induces callose formation at the sieve plate, reducing phloem transport upstream of the sieve plate but not below it. Carbon export and storage are hindered by the phloem transport dysfunction in infected plants.

Additionally, we observed that transport speed was moderately correlated with the number of callose plugs, but poorly correlated with the callose deposit size and density (Table 1). A greater mean CLas ΔCt value did lead to larger, denser callose deposits, but the size and density of deposits did not greatly affect phloem translocation speeds. It is likely that once a callose formation has closed off a route within the phloem, a larger, denser deposit makes no further difference to the total transport capacity, though greater bacterial populations stimulate continued callose deposition.

The two rootstocks examined here did not affect CLas ΔCt values. However, our data revealed that rootstocks produced different reactions in the scion to CLas infection in terms of both callose formation and phloem transport speed (Figures 2 and 4). This concurs with other studies that found that although symptom severity does vary significantly among rootstocks, this does not have a strict positive correlation with CLas titer, especially if plants have been infected for longer than a year (Folimonova et al. 2009; Albrecht and Bowman 2012). Healthy Hamlin plants on X-639 rootstock are less likely to produce phloem callose than those on Cleopatra (Figure 4; Supplementary Table S4). This results in a faster phloem translocation rate in healthy X-639 than healthy Cleopatra (Figure 2). However, when infected, plants on Cleopatra produced less total callose that was less dense and had a faster phloem translocation rate than infected plants on X-639 (Figure 2, Figure 4). Though there is little data about the sensitivity to these two genotypes to HLB, in some cases, hybrids between Cleopatra and *Poncirus trifoliata* provided higher tolerant to HLB, compared to Cleopatra (Albrecht and Bowman 2012), but this does not seem to be the case for X-639. Collectively, CLas-infected trees on X-639 rootstocks produced more callose, had slower sugar translocation and export rates, and accumulated less sucrose in the roots, although relatively partitioning more carbon into the roots compared to healthy plants, compared to those on Cleopatra rootstocks.

The site on the plant stem did not have effect on CLas ΔCt value. This is corroborated by previous research that concluded that CLas titer was inconsistent in different sampling sites (Folimonova et al. 2009). However, there were some significant effects of stem site on callose deposit quantity and density, which also differed depending on the interaction between rootstock type and CLas status of the plant (Figure S3). These effects were complex and varied in both effect size and direction. More extensive study is needed to clarify these relationships. However, the results of the regression between ΔCt values, callose deposit counts, and phloem translocation speeds suggest the hypothesis that translocation speed is affected by the other two factors on a local level (Figure 5). Considering the results of de Schepper et al. (2013), indicating that physical interruption of phloem transport interrupts transport upstream, but not downstream of the blockage, the present results may be interpreted as showing that CLas induces interruption of phloem function all along the trunk pathway. Figure 6 illustrates our proposed model for phloem transport disruption caused by CLas-induced callose formation.

The accumulation of sucrose and starch and allocation of ^14^C label further corroborate the interpretation of CLas infection as causing a transport limitation. Leaf export rates maintained the same pattern as transport, despite no observation of the accumulation of foliar starch. Root sucrose, however, was affected, with the greatest impact occurring in structural roots of X-639. This reduction in root sucrose of X-639 corroborates the reduced foliar sucrose export (Figure 2). This further suggests reduced mobilization of sucrose from structural roots, which act as a storage tissue (Goldschmidt & Koch 2017), to the fibrous roots that are considered sink tissues requiring the carbon for active growth. While sucrose content in fibrous roots show no statistical difference, fibrous roots of CLas-infected trees show numerically lower sucrose levels. This implies that root morphology and architecture could be negatively affected by the CLas infection in X-639 as suggested by the accumulation and mobilization of carbohydrates within the root system, a result corroborated by root observations of Johnson et al. (2013).

The starch results further strengthen this interpretation, with CLas infection causing a strong reduction of starch content in fibrous roots but not in structural roots (Supplemental Figure S2) in X-639. Together, because no change in starch level (Supplemental Figure S2) but reduced sucrose (Figure 3) were observed in structural roots of infected X-639, and possibly reduced sucrose in fibrous roots, the flux of carbon to storage as starch appears to be prioritized in infected X-639 roots over movement to fibrous roots. The relative activity of the ^14^C label in the roots is also consistent with this pattern, with nearly 6x greater accumulation of the label in roots of X-639 due to infection. This pattern is consistent with transport limitation along the whole transport path because such a limitation would greatly reduce the loss of labeled carbon in roots through respiration and the dilution of the label signal through new root growth, while affecting delivery to the intermediate sink to a lesser degree.

This transport impact was implicitly hypothesized in studies that observed reduced leaf carbohydrate export along with callose deposition (Koh et al. 2012), however, other studies have hypothesized tissue-specific metabolic effects, either primarily impacting leaves (Fan et al. 2010; Rao et al. 2018) or roots (Johnson et al. 2014a,b). The results of the present study are consistent with previous studies in finding impacts of CLas infection on carbon transport, leaf, and root sugar metabolism. By quantifying transport and carbohydrate distribution along the transport path, our data corroborate the previously observed symptoms, which are consistent with phloem transport limitation in CLas-infected trees (Figure 6).

Our results have verified a connection between the presence of CLas, the accumulation of callose in the stem phloem, and phloem transport limitation. The effect of callose formation and CLas ΔCt value on phloem speed is clear, but the contribution of rootstock type and the stem site appears more complex and requires further studies for clarification. Moreover, the precise mechanism of these connections is still unknown. It is likely that CLas both directly and indirectly causes phloem transport limitation, and this is the first quantitative demonstration of transport limitation caused by a plant microbial infection. This work provides a model testing system that can be used to directly measure plant host responses to phloem-limited pathogens and can serve as a foundation to further explore aspects of the CLas-*Citrus* pathosystem and the effects of other phloem-limited pathogens in the future.

## Acknowledgments

Funds to purchase equipment used for translocation speed measurement in this work were received from the University of Florida Institute of Food and Agricultural Sciences Equipment Seed Fund program. This work was supported by the University of Florida, Institute of Food and Agricultural Sciences Early Career Seed Grant (No. 00127818) and start-up funds.

## Author Contributions

CV, AL, and MD conceived, designed, and provided infrastructure and funding for experiments. MP performed ^14^C experiments. JPS performed starch and sucrose quantification. SW performed callose imagery and qPCR data collection. SW analyzed callose, PCR, and transport speed data with input from CV and AL. CV analyzed sucrose, starch, allocation, and leaf export data. Manuscript was written by SW with revisions and input from JPS, AL, and CV. All authors approved the final version of the manuscript.

