## Supplemental Figures for "Phloem transport limitation in Huanglongbing affected sweet orange is dependent on phloem-limited bacteria and callose"

**Supplemental Figures (Welker et al.):**

**Supplemental Figure S1.** Foliar sucrose concentrations (mM glucose equivalents) of leaves of Hamlin sweet orange (*Citrus sinensis*) scion on Cleopatra mandarin (*C. reticulata*) or X-639 (*C. reticulata* x *Poncirus trifoliata*) infected or uninfected by *Candidatus* Liberibacter asiaticus. Although the rootstock x CLas status interaction was *P*=0.04, no significant differences were detected at *P*<0.05 by Fisher’s LSD.

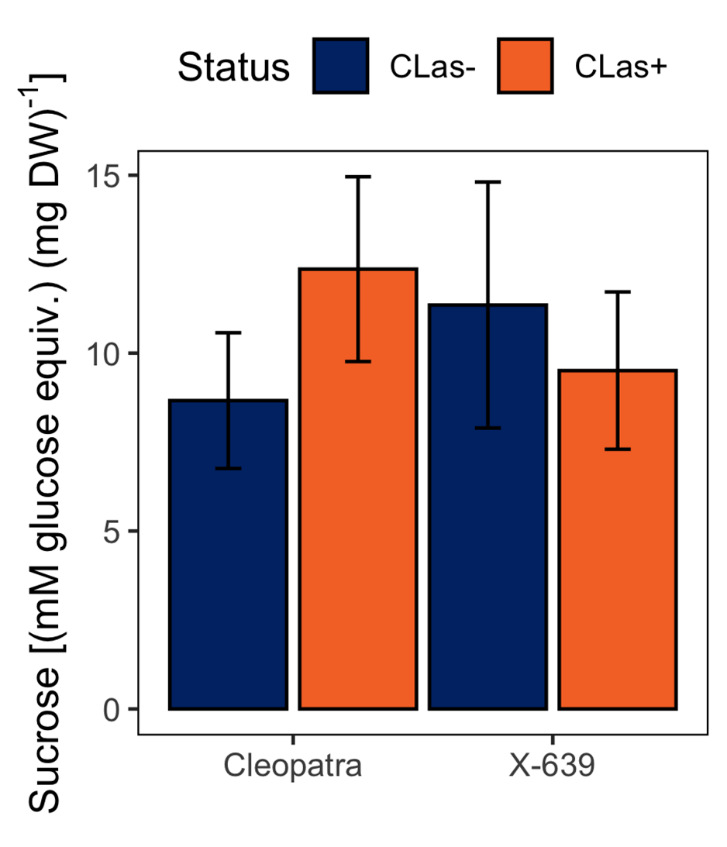

**Supplemental Figure S2.** Starch concentrations (mM glucose equivalents) of fibrous and structural roots of Cleopatra mandarin (*Citrus reticulata*) or X-639 (*C. reticulata* x *Poncirus trifoliata*) infected or uninfected by *Candidatus* Liberibacter asiaticus, all with Hamlin sweet orange (*C. sinensis*) scion. Bars show means, and error bars show standard error. Bars labeled with different letters are significantly different at P<0.05 by Fisher’s LSD.

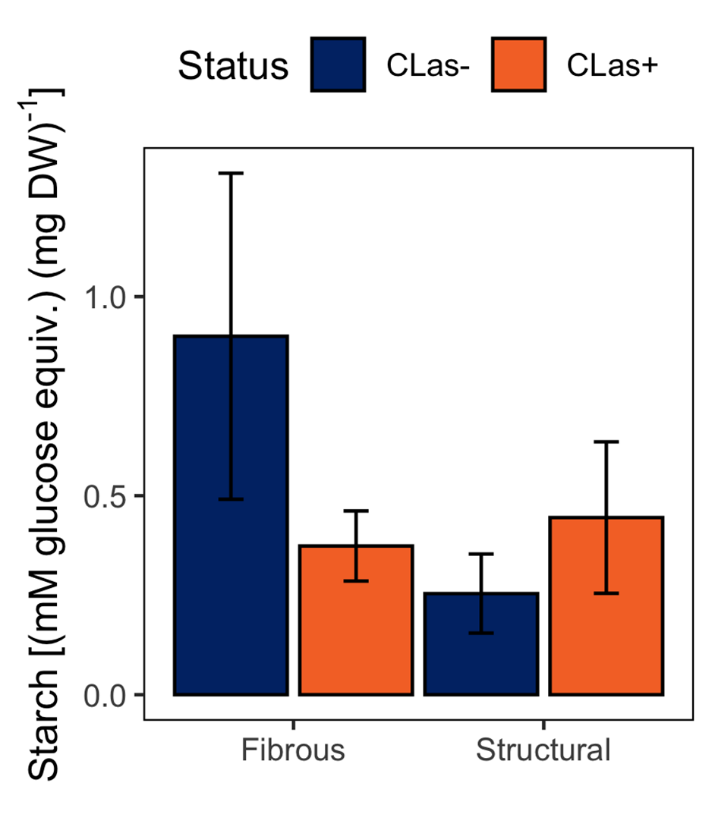

**Supplementary Figure S3: (**Top) Mean number of callose deposits detected per 10x field in confocal micrographs of citrus phloem. One 10x field image captures an area of phloem tissue which is approximately 1.16 mm^2^. Error bars represent standard error, number over each error bar represents n. All contrasts are significantly different at *P*= < 0.05 by Fisher’s LSD. (Bottom) Mean integrated density of callose deposits measured in the phloem. Integrated density is the product of the callose deposit area and its fluorescence brightness in relative fluorescence units (RFU), which approximates size and density. Error bars represent standard error, the number above each bar represents n. The large differences in n of integrated density measurements are the result of the wide variation of mean callose deposit counts per image exhibited between groups. The difference between the means of the CLas- and CLas+, X-639 and Cleo groups are all significant at *P* < 0.05 according to analysis of variance results. The difference between the mean of site 3 and the means of other sites is significantly different at *P*< 0.05 according to Bonferroni’s adjusted LSD.

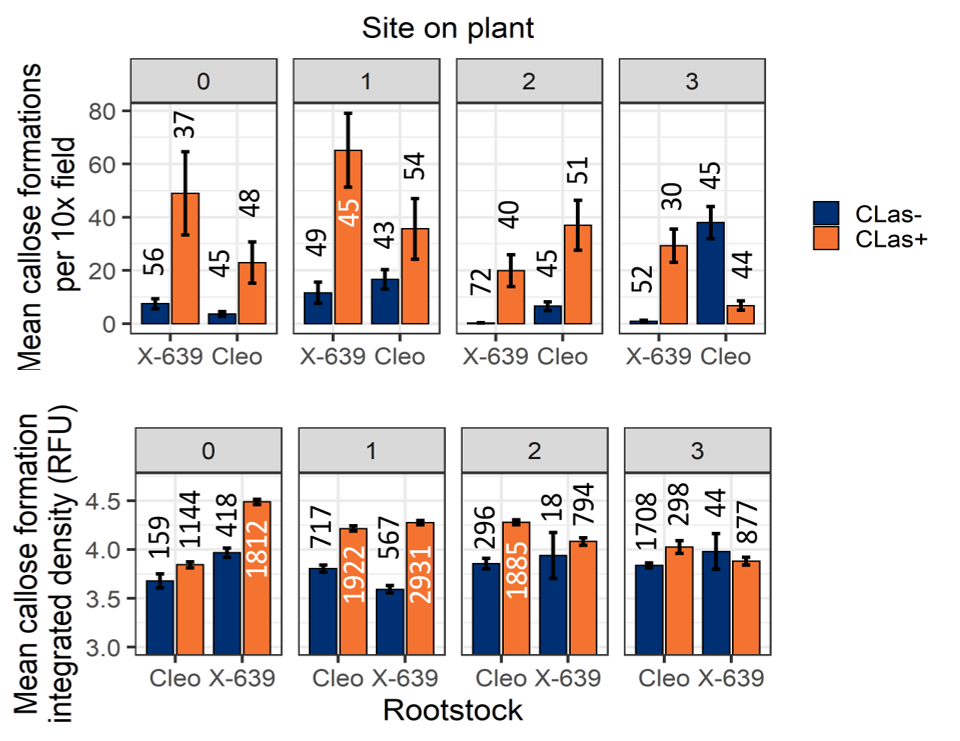
