## Supplemental Tables for "Phloem transport limitation in Huanglongbing affected sweet orange is dependent on phloem-limited bacteria and callose"

**Supplemental Tables (Welker et al.)**

| **Supplemental Table S1**  Analysis of variance of ∆Ct values from qPCR to amplify CLas genetic material in tissue samples from healthy and infected Hamlin/Cleopatra and Hamlin/X-639 plants. | | | |
| --- | --- | --- | --- |
| Effect | Degrees of Freedom | *F*-value | *P*-value |
| Rootstock | 1 | 1.67 | 0.206 |
| CLas | 1 | 54.49 | **<0.001** |
| Site on plant^1^ | 5 | 0.32 | 0. 898 |
| Rootstock x CLas | 1 | 0.05 | 0.822 |
| Rootstock x site on plant | 5 | 0.41 | 0.836 |
| CLas x site on plant | 5 | 0.39 | 0.853 |
| Rootstock x CLas x site on plant | 5 | 0.09 | 0.993 |

^1^Tissue samples were collected from the same 3 sites where phloem translocation speeds were measured, in addition to the leaf petiole.

| **Supplemental Table S2**  Analysis of variance of phloem translocation speeds assessed at three sites in the stems of CLas- or CLas+ Hamlin sweet orange on either Cleopatra or X-639 rootstock. | | | |
| --- | --- | --- | --- |
| Effect | Degrees of Freedom | *F*-value | *P*-value |
| Rootstock | 1 | <0.001 | 0.984 |
| CLas | 1 | 5.76 | **0.026** |
| Site on plant | 1 | 0.90 | 0.354 |
| Rootstock x CLas | 1 | 5.81 | **0.026** |
| Rootstock x site on plant | 1 | 0.86 | 0.364 |
| CLas x site on plant | 1 | 1.83 | 0.191 |
| Rootstock x CLas x site on plant | 1 | 2.28 | 0.147 |

**Supplementary Table S3:**

Analyses of variance of sucrose concentrations, starch concentrations, and relative 14C sink activity of leaf, trunk cortical tissues (bark), and root tissues (both structural root cortical tissues and fibrous roots with < 2 mm diameter) of Cleopatra mandarin (*Citrus reticulata*) or X-639 (*C. reticulata* x *Poncirus trifoliata*) infected or uninfected by *Candidatus* Liberibacter asiaticus, all with Hamlin sweet orange (*C. sinensis*) scion. Analyses were conducted separately for different tissue types. For leaves locations were distal and basal portions of the canopy using non-labeled leaves. For bark locations were 3 segments below the canopy relating to translocation speed measurements gathered in the same study. For roots locations were structural and fibrous roots. Values with P<0.1 are indicated in bold for ease of reading.

| Sucrose |  |  |  |  |
| --- | --- | --- | --- | --- |
| **Tissue type** | **Effect** | **DF** | **F-value** | **P-value** |
| Leaf | HLB | 1 | 0.28 | 0.61 |
|  | Rootstock | 1 | 0.00 | 0.97 |
|  | HLB x Rootstock | 1 | 6.00 | **0.04** |
|  | Location | 2 | 1.55 | 0.28 |
|  | HLB x Location | 2 | 0.51 | 0.62 |
|  | Rootstock x Location | 2 | 0.46 | 0.65 |
|  | HLB x Roostock x Location | 2 | 1.96 | 0.21 |
|  | Residuals | 7 |  |  |
| Bark | HLB | 1 | 1.03 | 0.33 |
|  | Rootstock | 1 | 0.08 | 0.79 |
|  | HLB x Rootstock | 1 | 0.00 | 0.95 |
|  | Location | 2 | 3.31 | **0.08** |
|  | HLB x Location | 2 | 0.10 | 0.91 |
|  | Rootstock x Location | 2 | 0.85 | 0.46 |
|  | HLB x Roostock x Location | 2 | 0.42 | 0.67 |
|  | Residuals | 10 |  |  |
| Roots | HLB | 1 | 8.09 | **0.01** |
|  | Rootstock | 1 | 2.83 | 0.12 |
|  | HLB x Rootstock | 1 | 1.13 | 0.31 |
|  | Location | 2 | 6.73 | **0.01** |
|  | HLB x Location | 2 | 0.08 | 0.92 |
|  | Rootstock x Location | 2 | 2.04 | 0.17 |
|  | HLB x Roostock x Location | 2 | 4.25 | **0.04** |
|  | Residuals | 12 |  |  |

| Starch |  |  |  |  |
| --- | --- | --- | --- | --- |
| **Tissue type** | **Effect** | **DF** | **F-value** | **P-value** |
| Leaf | HLB | 1 | 0.66 | 0.44 |
|  | Rootstock | 1 | 0.07 | 0.80 |
|  | HLB x Rootstock | 1 | 0.82 | 0.40 |
|  | Location | 2 | 1.43 | 0.30 |
|  | HLB x Location | 2 | 0.43 | 0.66 |
|  | Rootstock x Location | 2 | 1.93 | 0.22 |
|  | HLB x Roostock x Location | 2 | 2.18 | 0.18 |
|  | Residuals | 7 |  |  |
| Bark | HLB | 1 | 1.06 | 0.33 |
|  | Rootstock | 1 | 0.00 | 0.96 |
|  | HLB x Rootstock | 1 | 2.34 | 0.16 |
|  | Location | 2 | 3.21 | 0.08 |
|  | HLB x Location | 2 | 0.01 | 0.99 |
|  | Rootstock x Location | 2 | 1.90 | 0.20 |
|  | HLB x Roostock x Location | 2 | 0.05 | 0.95 |
|  | Residuals | 10 |  |  |
| Roots | HLB | 1 | 1.85 | 0.20 |
|  | Rootstock | 1 | 2.07 | 0.18 |
|  | HLB x Rootstock | 1 | 0.61 | 0.45 |
|  | Location | 2 | 1.35 | 0.30 |
|  | HLB x Location | 2 | 3.40 | **0.067** |
|  | Rootstock x Location | 2 | 0.86 | 0.45 |
|  | HLB x Roostock x Location | 2 | 0.04 | 0.96 |
|  | Residuals | 12 |  |  |

| 14C allocation | |  |  |  |
| --- | --- | --- | --- | --- |
| **Tissue type** | **Effect** | **DF** | **F-value** | **P-value** |
| Leaf | HLB | 1 | 3.44 | 0.11 |
|  | Rootstock | 1 | 1.91 | 0.21 |
|  | HLB x Rootstock | 1 | 0.25 | 0.63 |
|  | Location | 3 | 0.82 | 0.52 |
|  | HLB x Location | 2 | 0.59 | 0.58 |
|  | Rootstock x Location | 2 | 0.22 | 0.81 |
|  | HLB x Roostock x Location | 2 | 0.12 | 0.74 |
|  | Residuals | 7 |  |  |
| Bark | HLB | 1 | 0.01 | 0.92 |
|  | Rootstock | 1 | 3.97 | 0.07 |
|  | HLB x Rootstock | 1 | 3.85 | 0.08 |
|  | Location | 3 | 5.15 | **0.02** |
|  | HLB x Location | 3 | 1.47 | 0.28 |
|  | Rootstock x Location | 3 | 3.57 | **0.05** |
|  | HLB x Roostock x Location | 2 | 2.39 | 0.14 |
|  | Residuals | 11 |  |  |
| Roots | HLB | 1 | 0.18 | 0.68 |
|  | Rootstock | 1 | 0.46 | 0.51 |
|  | HLB x Rootstock | 1 | 8.39 | **0.01** |
|  | Location | 2 | 4.33 | **0.04** |
|  | HLB x Location | 2 | 1.24 | 0.32 |
|  | Rootstock x Location | 2 | 1.40 | 0.28 |
|  | HLB x Roostock x Location | 2 | 1.21 | 0.33 |
|  | Residuals | 13 |  |  |

| **Supplemental Table S4**  Significant results of the zero-altered negative binomial (ZANB) regression of callose deposit counts in the stem phloem of healthy and infected Hamlin/Cleopatra and Hamlin/X-639 plants^1^. | | | | |
| --- | --- | --- | --- | --- |
| Hurdle portion^2^ | | | | |
| Effect | Estimate^3^ | Standard Error | z-value | *P*-value |
| Rootstock is Cleo | 1.54 | 0.62 | 0.62 | **0.013** |
| Plant is CLas+ | 0.09 | 0.68 | 3.62 | **<0.001** |
| Site is 2 | -0.24 | 0.09 | -3.09 | **0.002** |
| Site is 3 | -0.24 | 0.09 | -2.77 | **0.006** |
| Rootstock is Cleo and site is 0 | -0.40 | 0.11 | -3.68 | **<0.001** |
| Plant is CLas+ and site is 0 | -0.39 | 0.12 | -3.26 | **0.001** |
| Plant is CLas+ and site is 3 | -0.77 | 0.26 | -2.98 | **0.003** |
| Rootstock is Cleo and plant is CLas+ and site is 0 | 0.68 | 0.16 | 4.33 | **<0.001** |
| Rootstock is Cleo and plant is CLas+ and site is 3 | 1.00 | 0.28 | 3.45 | **<0.001** |
| Count portion^4^ | | | | |
| Effect | Estimate | Standard Error | z-value | P-value |
| Site is 0 | -0.27 | 0.08 | -3.56 | **<0.001** |
| Site is 2 | -0.72 | 0.16 | -4.60 | **<0.001** |
| Site is 3 | -0.24 | 0.09 | -2.77 | **0.005** |
| Rootstock is Cleo and plant is CLas+ | -3.15 | 1.00 | -3.17 | **0.002** |
| Rootstock is Cleo and site is 2 | 0.48 | 0.17 | 2.85 | **0.004** |
| Rootstock is Cleo and site is 3 | 0.46 | 0.14 | 3.21 | **0.001** |
| Plant is CLas+ and site is 3 | -0.77 | 0.26 | -2.98 | **0.003** |

**^1^** All combinations of all the factor levels were tested. Only significant results were included here for brevity. **^2^**Hurdle or binomial portion of the model estimated the odds of whether callose will form at the sieve pore or not. ^3^Estimates indicate the effect size of the factor, while the sign of the estimate indicates the effect direction; a negative sign indicates the factor made callose deposits less likely while a positive sign indicates that callose formation was more likely. ^4^Count portion of the model estimated how many callose deposits will accumulate if the callose deposit count is non-zero.

| **Supplemental Table S5**  Analysis of variance of mean integrated density measurements^1^ of callose deposits in the stem phloem of healthy and infected Hamlin/Cleopatra and Hamlin/X-639 plants. | | | |
| --- | --- | --- | --- |
| Effect | Degrees of Freedom | F-value | P-value |
| CLas | 1 | 390.24 | **<0.001** |
| Rootstock | 1 | 17.23 | **<0.001** |
| Site on plant | 1 | 4.21 | **0.040** |
| Rootstock x CLas | 1 | 18.16 | **<0.001** |
| Rootstock x site on plant | 1 | 266.42 | **<0.001** |
| CLas x site on plant | 1 | 7.67 | **0.006** |
| Rootstock x CLas x site on plant | 1 | 104.52 | **<0.001** |

^1^ Measurements were Box-Cox transformed before computing means and ANOVA testing.
